# Longitudinal sequencing reveals polygenic and epistatic nature of genomic response to selection

**DOI:** 10.1101/2024.07.22.604612

**Authors:** Simon K.G. Forsberg, Diogo Melo, Scott Wolf, Jennifer K. Grenier, Minjia Tang, Lucas P. Henry, Luisa F. Pallares, Andrew G. Clark, Julien Ayroles

## Abstract

Evolutionary adaptation to new environments likely results from a combination of selective sweeps and polygenic shifts, depending on the genetic architecture of traits under selection. While selective sweeps have been widely studied, polygenic responses are considered more prevalent but challenging to quantify. The infinitesimal model makes explicit the hypothesis about the dynamics of changes in allele frequencies under selection, where only allelic effect sizes, frequencies, linkage, and gametic disequilibrium matter. Departures from this, like long-range correlations of allele frequency changes, could be a signal of epistasis in polygenic response. We performed an *Evolve & Resequence* experiment in *Drosophila melanogaster* exposing flies to a high-sugar diet as a source of environmental stress for over 100 generations. We tracked allele frequency changes in >3000 individually sequenced flies as well as population pools and searched for loci under selection by identifying sites with allele frequency trajectories that differentiated selection regimes consistently across replicates. We estimate that at least 4% of the genome was under positive selection, the result of a highly polygenic response. Most of this response was seen as small but consistent allele frequency changes over time, and there were only a few large allele-frequency changes (selective sweeps). We then searched for signatures of selection on pairwise combinations of alleles in the new environment and found several strong signals of putative epistatic interactions across unlinked loci that were consistent across selected populations. Finally, we measured differentially expressed genes (DEGs) across treatments and show that DEGs are enriched for selected SNPs, suggesting a regulatory basis for the selective response. Our results suggest that epistatic contributions to polygenic selective response are common and lead to detectable signatures.

## INTRODUCTION

Genetic changes underlying evolutionary response to a new environment can differ depending on the genetic architecture of the traits under selection. For traits with simple genetic architectures, controlled by few genes with large allelic effect sizes, we would expect to see selective sweeps, in which a positively selected allele rapidly goes to fixation and leaves a detectable signature in the surrounding genomic regions (Pavlidis & Alachiotis, 2017; Smith & Haigh, 1974). Interest in finding the causal allele responsible for a phenotype has caused a bias for this type of selective response in the literature (Pritchard & Di Rienzo, 2010), and we have cataloged several positively selected genes in humans and other species (Feder et al., 2016; Garud et al., 2015; Ihle et al., 2006). In contrast, for traits that are controlled by many genes, quantitative genetic theory predicts that selection response is generated by more subtle changes across the allele frequencies of many loci (Chevin & Hospital, 2008; Jain & Stephan, 2017), and indeed it has become clear that this is a more common form of adaptive response than hard sweeps (Barghi et al., 2020; Pritchard et al., 2010).

While much of the experimental effort has been focused on determining how many genomic regions respond to selection, there is more to genetic architecture than the number of loci affecting a trait (Hansen, 2006). Gene-by-gene (epistatic) and gene-by-environment interactions both contribute to genetic variation and can be leveraged in response to selection. The extent to which epistasis is important for polygenic response is still an open question (Crow, 2010; Csilléry et al., 2018; Hansen, 2013; Le Rouzic, 2014; Phillips, 2008; Weinreich et al., 2013). Given the possibility that epistasis contributes appreciably to adaptation, we argue that polygenic selection response should result in two observable patterns when epistasis is present: i) a correlation between the allele frequencies at interacting loci—i.e. change in allele frequency at one locus is accompanied by corresponding changes at the (potentially unlinked) interacting loci (Csilléry et al., 2018), and ii) the emergence of gametic disequilibrium in adapted populations—as allelic combinations are selected for or against—resulting in deviations from two-locus Hardy-Weinberg proportions between pairs of unlinked loci (Boyrie et al., 2021

Identifying these signatures of polygenic and epistatic response to selection is a challenging problem and Evolve and Resequence (E&R) experiments have emerged as a natural and powerful tool for investigating these questions (Barghi et al., 2020). By exposing replicate populations to a stressful treatment condition and keeping them in this environment for several generations we are able to track the resulting changes in genetic composition due to selection. By tracking allele frequencies through time in both selected and control populations, we can distinguish the effects related to adaptation to laboratory conditions and the effects of the selective stress to which treatment populations are exposed. The environmental change caused by the exposure to stress also opens the possibility for new gene combinations to come under selection (Das et al., 2020; Ogbunugafor, 2022), allowing us to search for the signatures of selection on epistatic combinations that are advantageous in the new environment.

Alterations in the diet is likely to be a ubiquitous source of environmental stress for animal populations. When exposed to varying levels of dietary sugar, *Drosophila melanogaster* individuals display complex metabolic and behavioral responses (Chng et al., 2017; McKenzie & McKechnie, 1979). Exposure to high sugar can lead to changes in the absorption and metabolism of sugars (Hickey & Benkel, 1982; Zinke et al., 2002), along with altered foraging and feeding behaviors (Dus et al., 2015; Lim et al., 2014). These metabolic responses are mediated by neuronal and endocrine signaling networks in the head, which interact with and cause coordinated responses in the corpora cardiaca and fat bodies in the thorax and abdomen, resulting in an organism-wise response (Chng et al., 2017). High sugar also causes differential gene expression, leading to up regulation of digestive enzymes and other genes involved in lipid metabolism (Chng et al., 2014; Mattila et al., 2015). Flies exposed to chronic high sugar can develop several pathologies, like obesity, diabetes-like responses, cardiomyopathy, shorter life span, and tumor growth (Birse et al., 2010; Musselman et al., 2011; Na et al., 2013; Pallares, Lea, et al., 2020). This suggests that high-sugar stress is likely to cause generalized responses in several interacting gene networks and should lead to strong selective pressures.

Here, we performed an E&R experiment where three replicate populations of Drosophila melanogaster were exposed to a stressful environment in the form of a high level of dietary sugar, while another three replicate populations were maintained on a control diet. All six populations were derived from the same base population. Whole genome sequencing was performed on flies from all six populations at generations 1, 11, 25, and 100, giving a total of almost 3000 sequenced individuals (fig. 1 A). Using this time series genomic data, we identify two directions of allele frequency change. The largest driver was shared across selected and control populations, suggesting a shared lab environment selection. The second-largest driver of genetic change contrasts control and selected populations, and so is linked to the selection regime. We estimate that at least 4% of the genome was under positive selection due to high-sugar stress. Most observed changes in allele frequency are however relatively modest, and using the individual sequence data to estimate haplotypes, we show how most of the selected loci do not show archetypal signals of selective sweeps after 100 generations. These results point towards a highly polygenic selection response, in line with theoretical expectations from quantitative genetics theory. We then measure differentially expressed (DE) genes across selected and control lines after selection and show that DE genes are highly enriched for selected SNPs, pointing to regulatory divergence as a mechanism for the effects of the polygenic response to selection. We also quantify correlations in allele frequency between pairs of selected loci over time, as well as gametic disequilibrium after 100 generations of adaptation, and show that several alleles show correlations and gametic disequilibrium across unlinked loci, suggesting that epistatic interactions participated in the response to selection. We confirm this last point by using Wright-Fisher simulations, showing that the correlations and the gametic disequilibrium we observe are unlikely to appear in the absence of epistatic interactions. While we lack a clear phenotype to directly study the effect of epistatic variation on selection response, our results suggest that epistatic contributions to polygenic response to selection are common and lead to detectable genomic signatures.

**Figure 1:**
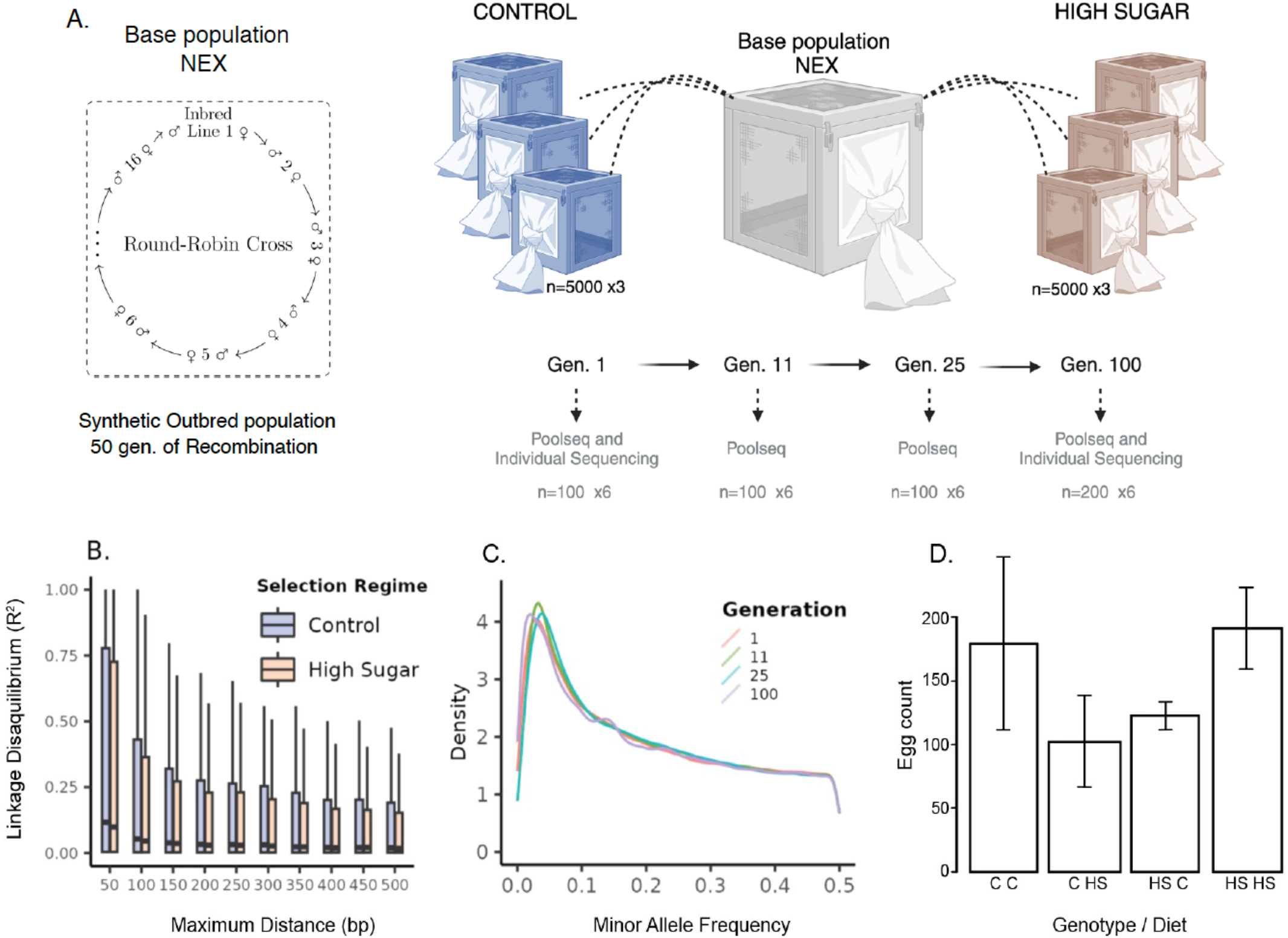
Selection experimental design. **A**. Scheme of the experimental design. A synthetic outbred population was created by a round-robin cross of 16 lines from the Netherlands. This population (NEX) was kept as an outbred population for over 50 generations before the start of the selection experiment. Starting from NEX, 3 control (control) and 3 treatment (hs) populations of around 5000 individuals were kept for 100 generations. Samples of one hundred individuals were taken at generations 1, 11, 25 and 100 for allele frequency tracking. **B**. Linkage disequilibrium (LD) decay across hs and control populations at generation 100. NEX derived populations have very low levels of LD. **C**. Minor allele frequency across generations. **D**. Egg-lay measurements after selection showing the adaptive response to the high-sugar environment in hs populations Control flies on Control food (C C), Control flies on High Sugar food (C HS), High Sugar selected flies on Control food (HS C), High Sugar selected flies on High Sugar food (HS HS).

## Results

### Polygenic Selection Response

To study the effect of long-term selection on a stressful environment, we kept three replicate populations of flies under high-sugar stress and three under control conditions for 100 generations. To assess if selected populations had adapted to the stressful high-sugar environment, we performed a factorial egg-lay experiment, measuring the fecundity of both control and high-sugar (hs) populations in the control and high-sugar diet. Both populations show higher fecundity in the corresponding diet (fig. 1 D), indicating successful adaptation. We collected data from 100 individuals at four time points and obtained both allele frequencies from Pool-seq and genotypes from individual sequencing (fig. 1 A and C). This time series genomic data allowed us to analyze the changes in allele frequency and identify the largest drivers of genetic change in response to the stressful environment. After quality control (see Methods), we obtained allele frequency estimates for ∼1.76M SNPs, giving a total data collection of 4 time points x 3 replicate populations x 2 treatments x 1.76M SNPs genotype calls. To identify the main drivers of genetic change without any prior assumptions, we performed a Principal Component Analysis (PCA) of the allele frequencies across the entire selection experiment. The first two principal components (PC), explaining 17% and 13% of the variance, largely coincides with time and selection regime, respectively (figs. 2, 7). This unsupervised approach thus identified time and high-sugar selection as the two main drivers of genetic change genome wide. Surprisingly, the time dimension, captured by the first PC, explained slightly more variance than the selection regime, which was captured by the second PC. This shows that all six populations experience some common selection pressures, presumably related to lab environment. The second largest driver of genetic change is indeed exposure to high-sugar stress (fig. 2).

**Figure 2:**
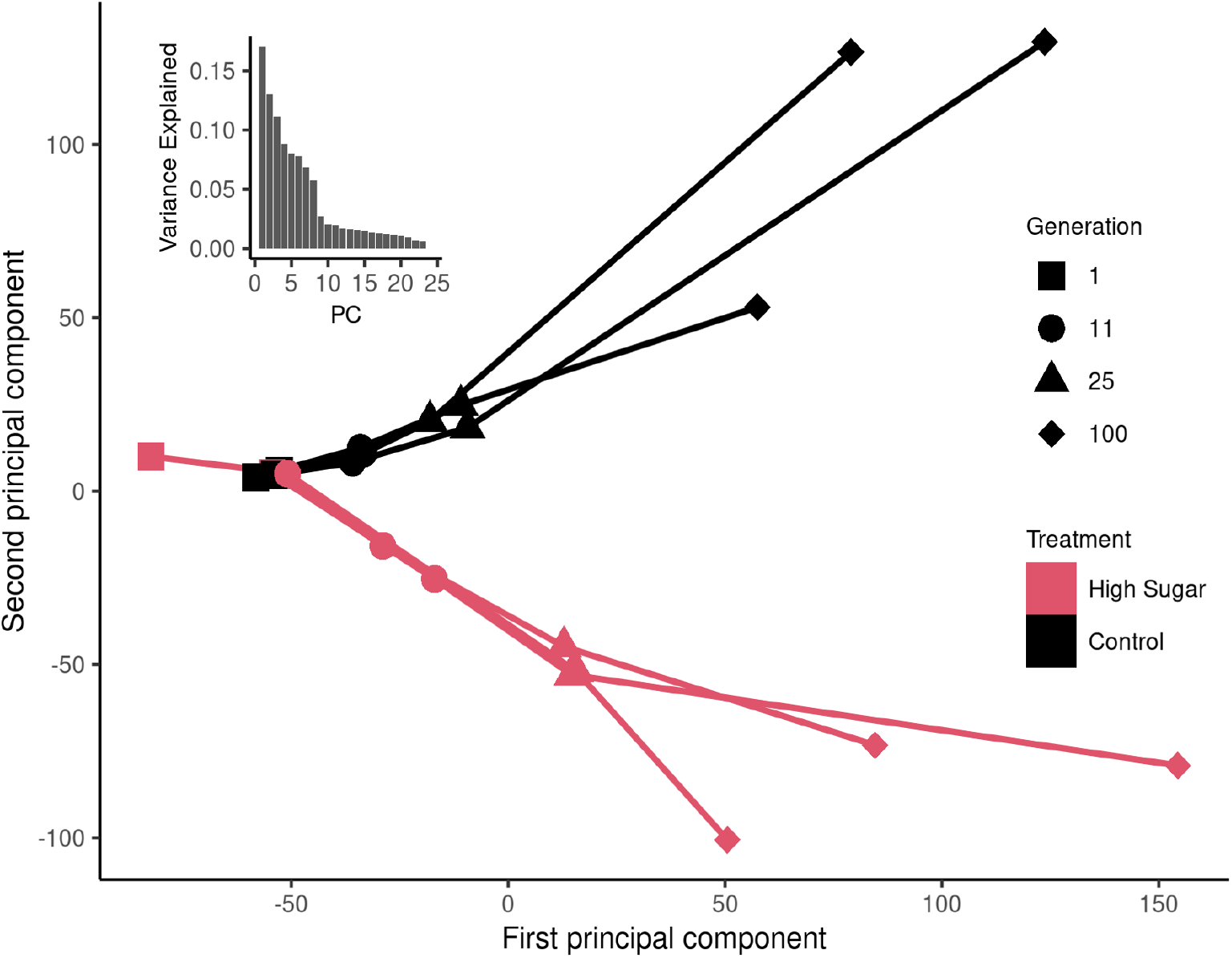
Principal components one (x-axis) and two (y-axis) from the PCA on the genome wide allele frequencies across the entire selection experiment. ***Each*** line corresponds to one of the six experimental populations, red indicating high-sugar treatment and black control, with symbols marking the mean scores for each population and time point. The variance explained by each of the first 23 principal components is shown in the inset.

### High-sugar selection on individual loci

To identify individual loci under selection, we fitted a univariate regression model for each SNP, incorporating allele frequencies across all time points, replicate populations, and selection regimes. This model identifies SNPs whose allele frequency changes in the same direction over time in all replicate populations. The time coefficient in the model captures changes that are similar across all six populations (fig. 3 A), and the time-by-selection-regime coefficient captures changes that are unique to one selection treatment (fig. 3 B and C). The p-values of the time coefficient were highly correlated with SNP loadings onto PC1 (cor = 0.59, p < 10-16, sup fig. 1), whereas the the p-values of the time-by-regime coefficient were highly correlated with SNP loadings onto PC2 (cor = 0.68, p < 10-16, sup fig. 1), consistent with the first two PCs capturing time and selection regime.

**Figure 3:**
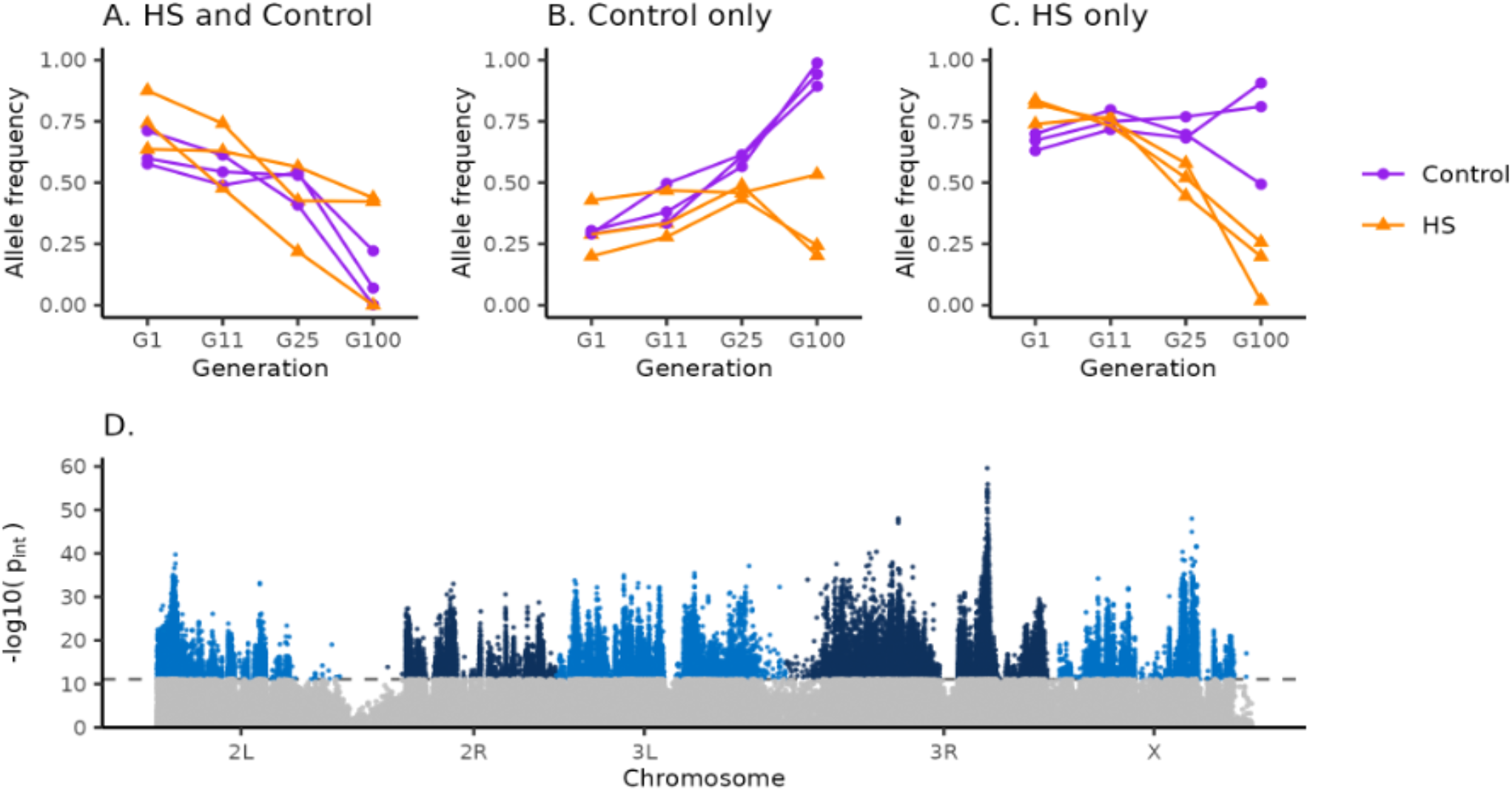
Results from the per SNP regression model. Panels A-C show possible patterns of relevant and consistent allele frequency change across the six populations. We chose SNPs with strong signals for illustration, but many significant SNPs show more subtle allele frequency changes. Plotted are allele frequencies trajectories of SNPs with significant linear trends under the specified model. **A**. Consistent change in control and hs, **B**. consistent change in control only, **C**. hs and control differ. Both the examples shown in B. and C. would lead to a significant interaction term between time and treatment, but we filter SNPs that change only in Control (like in panel B). **D**. Manhattan plot showing negative log10 transformed p-values from the regression analysis of allele frequency over time. The p-values correspond to the time-by-selection regime interaction coefficient in the model. A significant p-value indicates different trajectories in the two treatments. SNPs showing a selection response primarily in the control regime where excluded and are not shown.

Different SNPs showed very different allele frequency trajectories over time. Some respond similarly to selection in all replicate populations regardless of selection regime (fig. 3 A), while others respond in opposite directions (fig. 3 C) or in only one of the regimes (fig. 3 B). Our regression model allowed us to distinguish these different scenarios and, for what follows, we focus on the selection signatures that are unique to the high-sugar selection regime. The Manhattan profile in fig. 2 D, showing the timeby-regime p-values, suggests a polygenic selection response. This is in line with the observation that time and selection regime are the two main drivers of genetic change genome-wide (fig. 2).

In order to further relate the locus-specific results (fig. 3) to the genome-wide signal quantified by the PCA (fig. 2), we repeat the PCA after excluding all SNPs with a regression p-value below a given threshold, effectively removing the SNPs that are associated with the selection regime. Changing the significance threshold allowed us to evaluate the effects of the filtered SNPs on the PCA. When using a very conservative threshold, excluding only the most strongly selected SNPs, the results from the PCA remained largely unchanged, showing that the PCA signal is not driven by a few loci under very strong selection (sup fig. 2 A). We used these changes in the PCA as a heuristic to pick a p-value threshold of 8×10-12, since PC2 did no longer distinguish the different selection regimes when excluding SNPs with a p-value below this threshold (sup fig. 2 C). SNPs passing this significance threshold are thus driving the majority of the selection response to highsugar stress that we observe in the PCA.

### What proportion of the genome is responding to selection?

Using this conservative threshold, ∼45k SNPs show a signature of positive selection that is unique to the high-sugar selection regime. Considering 200 bp around every selected SNP, corresponding to an average *r*^*2*^ of 0.2 (fig. 1 B), these SNPs span ∼5.6 Mb, or ∼4% of the mappable genome of *D. melanogaster*. Since the linkage disequilibrium (LD) around the selected loci is expected to be larger than the genome-wide average, we believe this to be a conservative estimate. The magnitudes of the allele frequency changes tend to be relatively small. Comparing generation 1 to generation 100, the mean change across all SNPs in the populations exposed to the high-sugar selection regime is 0.11, while the mean change among the selected SNPs is 0.25 (fig. 4). Among all the 1.76 M SNPs, only 4753 show a pattern where the minor allele at generation 1 has reached fixation at generation 100 in at least one of the populations in the high-sugar selection regime. Furthermore, many SNPs also display a delayed selection response, with the largest change in allele frequency after generation 25 (fig. 4). This is consistent with theoretical predictions for polygenic adaptation involving independent loci (Chevin & Hospital, 2008; Pavlidis et al., 2012), but could also be due to epistatic effects (Paixão & Barton, 2016).

**Figure 4:**
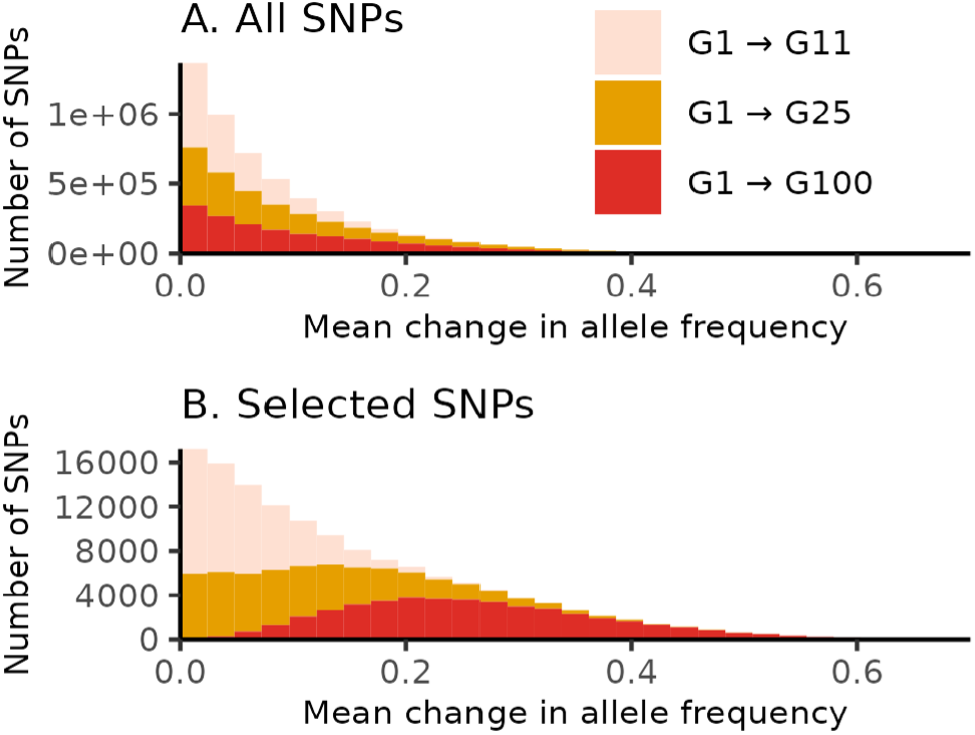
Histograms showing mean changes in allele frequency in the populations exposed the high-sugar selection regime, between generation 1 and 11, 1 and 25, and 1 and 100. Panel A includes all 1.76M SNPs and panel B includes the 45k SNPs that show a signature of positive selection unique to the high-sugar selection regime.

### Do the selected alleles show a detectable sweep signature?

Next, we ask if the identified selection signatures tend to coincide with the genomic footprint of selective sweeps. Using a core set of 20k high confidence SNPs, we estimated individual haplotypes at generation 100. These haplotypes were then used to calculate the integrated Haplotype Score (iHS) (Voight et al., 2006) in the HS populations. A large iHS indicates an extended haplotype associated with one allele at a given SNP, a pattern characteristic of a selective sweep. The estimated iHS and the p-values from our regression model showed a small but significant correlation (cor = 0.07, p = 2.1×10^−17^, sup fig. 3), indicating a tendency of longer haplotypes at the selected loci. The observed correlation is, however, very modest, showing that the loci indicated to be under selection using our time series data do not display a strong sweep-like pattern after 100 generations of adaptation. At a nominal significance threshold of p < 0.05, only 4.7% of the selected loci, as inferred from the regression analysis, also displayed a significant iHS. Taken together, these observations are all consistent with the polygenic view of adaptation through subtle shifts in allele frequency at a large number of loci, and with selection acting primarily on standing genetic variation rather than novel mutations.

### Effects of Polygenic Adaptation on Gene Expression

Much of the genetic variation for complex traits resides in gene-regulatory regions (Albert & Kruglyak, 2015). Selection on complex traits might then be expected to act largely on this regulatory variation, resulting in changes in gene expression. To characterize the effect of selection on gene expression, we performed a full reciprocal experiment where flies, adapted to either high-sugar or control selection regimes, were reared in either high-sugar or control conditions (fig. 5). This design allows us to account for short-term plastic changes due exposure to a different diet and the long-term effect of selection. For each one of the four experimental groups, we then performed RNA-seq separately on bodies and heads (n ∼ 40 per group, see Methods). After quality control, we obtained expression for 8397 genes from the body samples, and 8298 genes from the head samples.

**Figure 5:**
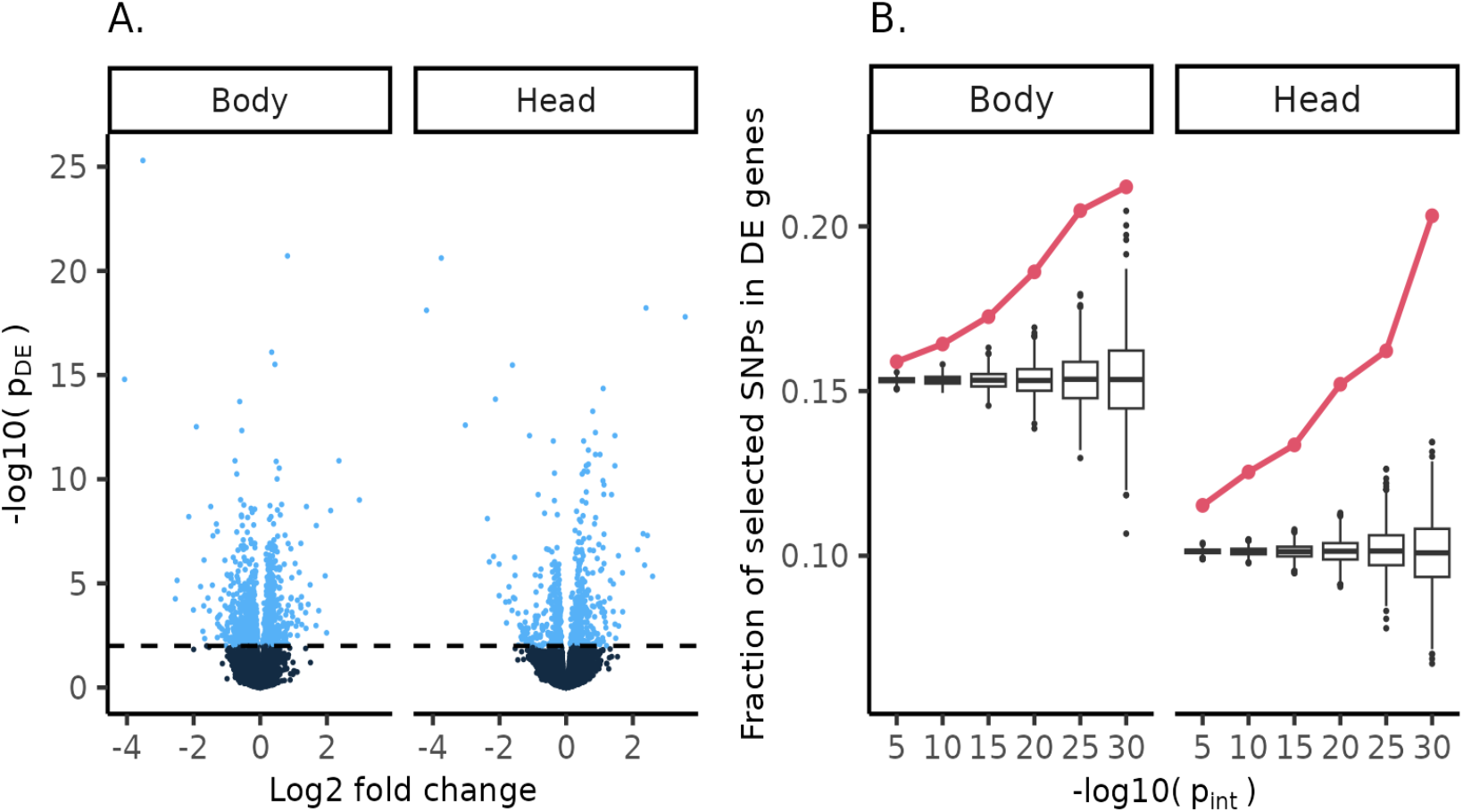
Differential expression after selection. **A**. Volcano plots showing differential gene expression between flies adapted to the high-sugar versus control selection regime, after controlling for the plastic effects related to each diet. Each point corresponds to one gene. Y-axis shows the negative log10 transformed p-value of the differential expression, and x-axis shows the log2 transformed fold change. The two panels correspond to expression in body and head tissue. **B**. Fraction of SNPs under positive selection in the high-sugar selection regime that coincide with a differentially expressed gene (y-axis), at different p-values for the selection signature (x-axis). Solid red lines show the observed fraction of SNPs overlapping DE genes and boxplots show the empirical null distributions obtained from permutations. The two panels correspond to expression in body and head tissue.

DE genes between flies adapted to the respective selection regimes was measured separately for head and body samples. At an FDR < 0.01, 1155 and 578 genes showed DE in body and head respectively. We went on to ask how many of these DE genes fall in regions with signatures of selection. In both body and head, we see an enrichment of selected SNPs among the genes showing DE (fig. 5 B). Starting at p-value of 10^−5^ for the selection term, this enrichment gets more pronounced with increasingly stringent selection p-values. The enrichment highly exceeds the expected random overlap between selection signals and DE genes, as estimated from a permutation test, indicating that adaptation appears to have acted on regulatory genetic variants.

### Epistasis across selected loci

Whether epistatic interactions contribute to long-term selection response is a contentious topic, but there are several possible mechanisms through which epistasis could play a role in polygenic response. For example, diminishing returns epistasis, where the rise in frequency of one allele decreases the fitness of alleles at other loci, which has been empirically observed (Kryazhimskiy et al., 2014), is implicit in stabilizing selection models describing population movement towards an optimum (Chevin & Hospital, 2008; Höllinger et al., 2019; Jain & Stephan, 2015). The access to allele frequency time-series and haplotype information allows us to look for two different signatures of epistatic contributions to the observed polygenic selection response. In the presence of epistasis for fitness, the trajectory of a given allele during the course of selection is not independent of the trajectories of alleles at other loci. Rather, the expected trajectory is a function of the trajectories of all other alleles that it interacts with (Paixão & Barton, 2016). This should result in two observable patterns: 1) a correlation between the allele frequencies at interacting loci, as changes in allele frequency at one locus is accompanied by corresponding changes at the interacting locus; 2) gametic disequilibrium in adapted populations, the selective removal of unfavorable allelic combinations should result in deviations from the 2-locus Hardy-Weinberg proportions expected for a pair of unlinked loci (Corbett-Detig et al., 2013).

To explore the expected genomic footprint of selection under fitness epistasis versus strict additivity in the current experiment, we performed a series of Wright-Fisher based simulations using the SLiM modeling framework (Haller & Messer, 2019). The simulations were set up to mimic the relevant aspects of the experiment, starting by creating neutral populations with about 3k segregating SNPs in mutation-drift equilibrium distributed along two unlinked chromosomes. Using these starting populations, we sampled 1000 segregating mutations to be a quantitative trait locus (QTL) contributing to the trait under truncation selection. We then simulated two scenarios: (1) an additive scenario, in which the value of the trait is only given by these additive QTLs, and (2) an epistatic scenario, where, in addition to the additive effects, we sampled 200 pairs of QTLs (one member of the pair in each chromosome) to have an additive-by-additive epistatic effect on the quantitative trait. After 100 generations of selection, we quantified both gametic disequilibrium and correlated changes in allele frequency. This was done for the same pairs of QTLs in both simulations, with the only difference being the presence of the epistatic interaction in one scenario. The gametic disequilibrium between unlinked loci was substantially higher in the presence of epistasis, as opposed to selection acting on a purely additive genetic architecture. Likewise, the correlation between allele frequency trajectories of QTL pairs in different chromosomes is only different from zero in the epistatic scenario (fig. 6 C).

**Figure 6:**
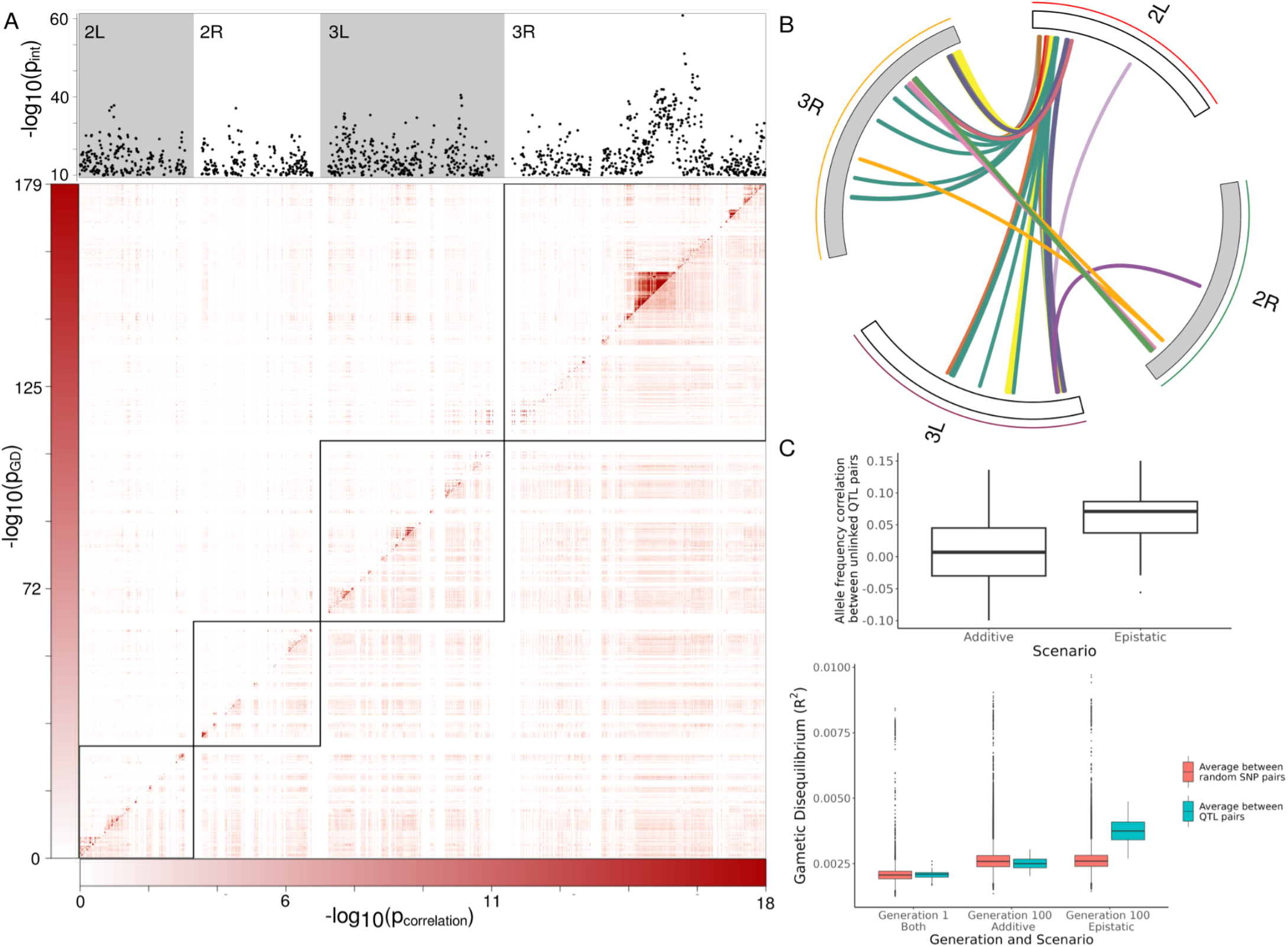
Signatures of epistasis in experimental data and simulations. **A**. Top panel shows negative log10 p-values (y-axis) from the regression analysis of allele frequency over time. The p-values correspond to the time-by-selection regime interaction coefficient in the model. SNP positions are scaled for visualization. Bottom panel shows a heatmap of pairwise SNP analyses performed in the populations exposed to the high-sugar selection regime. Negative log10 p-values of the gametic disequilibrium, given by a chi-square test, are shown above the diagonal. Negative log10 transformed p-values of the correlation in allele frequencies over time are shown below the diagonal. **B**. Locus pairs showing both genotype ratio distortions (chi-square test, p < 5.7×10^−8^) and correlated allele frequencies (correlation test, p < 0.001). The outer circle represents the chromosome arms, and each link represents a locus pair. Colors correspond to our LD clumping procedure, where links with the same color involve the same locus at one end, supported by multiple locally linked SNPs. **C**. Comparison of allele frequency correlations and gametic disequilibrium in the simulations, across additive and epistatic scenarios.

Next, we looked for the same kind of genomic footprint in our empirical data. For this analysis, we focused on a set of SNPs under strong selection unique to the high-sugar regime, indicated by a time-by-regime p < 8×10^−12^, and for which we had high enough coverage to confidently genotype a large number of individual flies, giving us a set of 1.3k SNPs. Having estimated the correlations in allele frequencies between these SNPs in the high-sugar regime, the resulting SNP×SNP correlation matrix gives a picture of the SNPs that move in unison through time in all replicate populations, and the ones that do not (fig. 6 A, lower triangle). Secondly, the SNP×SNP gametic disequilibrium matrix shows which SNP pairs display deviations from the proportions expected for unlinked loci in the populations exposed to the high-sugar selection regime (fig. 6 A, upper triangle).

Both allele frequency correlations and gametic disequilibrium were substantial between SNPs that are physically linked, as expected (fig. 6 A, elements near the diagonal). However, we also observe numerous examples of correlations between physically distant SNPs, showing that their allele frequencies change in a similar manner during the course of selection. This could be due to epistasis, but it could also be a consequence of similar but independent selection pressures acting on the two SNPs. However, we also identified multiple cases of gametic disequilibrium between physically distant SNPs, indicative of epistatic selection. Comparing gametic disequilibrium in the highsugar populations to the control populations, for the SNPs showing a signature of positive selection unique to the high-sugar regime, we observed a small but highly significant negative correlation (cor = -0.06, p < 10^−16^). This indicates that SNP pairs in gametic disequilibrium in one selection regime tend to segregate independently in the other. However, when doing the same comparison for the SNPs that display a similar signature of selection in both treatments, presumably due to lab environment adaptation, we observed a positive correlation (cor = 0.14, p < 10^−16^). This indicates that SNP pairs that are selected in both treatment also have more similar patterns of gametic disequilibrium. Taken together, this suggests that the observed gametic disequilibrium between physically distant SNPs is caused by epistatic selection, where allelic combinations that are under selection in the high-sugar regime tend to be neutral in the control regime, leading to gametic disequilibrium in the high-sugar populations but not the controls. On the other hand, allelic combinations that are conducive to fitness in both regimes are selected in a similar manner in all populations, creating a similar footprint of gametic disequilibrium in both high-sugar and control populations.

We identified 1413 SNP pairs in the HS populations where the two SNPs were located on different chromosomes, displayed gametic disequilibrium (chi-square test, p < 5.7×10^−8^), and correlated allele frequencies (correlation test, p < 0.001). To exclude the possibility that the observed gametic disequilibrium was due to population structure rather than epistasis, we also looked for evidence of gametic disequilibrium within each replicate population. Compared to jointly analyzing all replicate populations from the same selection regime, this analysis has lower power due to smaller sample size. Since, by design, no population structure exists within each replicate population, it cannot be the cause of gametic disequilibrium. Keeping only signals supported by multiple linked SNPs at “both ends”, and where at least two SNP pair gametic disequilibrium replicated within the replicate populations, we identify 11 pairwise epistatic selection signatures, each supported by between 5 and 63 SNP pairs (fig. 6 B).

## Discussion

Here, we track the allele frequency trajectories of 1.7 million SNPs in replicate populations of *D. melanogaster* exposed to either a control diet or a stressful high-sugar diet. Principal component analysis of the SNP allele frequencies in these population samples revealed a striking pattern: PC1 corresponded with generation time for both treatments, whereas PC2 cleanly separated the control from the high-sugar diet populations. By applying a regression model to the allele frequency trajectories between control and selected populations, we detected a large number of SNPs whose allele frequency changed consistently in the selected populations and not in the control population, suggesting that the exposure to high sugar drove the consistent response. Surprisingly, the lab environment, shared between control and selected populations, appears to be a stronger source of selection than the exposure to high sugar. The subtle changes in allele frequency and the large number of SNPs that show signatures of selection are suggestive of a massively polygenic selection response. This polygenic response appears to be grounded in regulatory changes, given that differentially expressed genes in the populations exposed to high sugar are enriched for SNPs under selection. We also look for signatures of interactions between selected loci, and find strong evidence for fitness epistasis in the form of gametic disequilibrium between physically unlinked loci and correlations between allele frequency trajectory of these same loci.

Highly polygenic genetic architectures have had a recent renaissance in the complex trait literature. Progressively more powerful genome-wide association studies (Pallares et al., 2023; Yengo et al., 2022), evidence from E&R experiments (Barghi et al., 2019; Burny et al., 2021) and gene-regulatory networks (Võsa et al., 2021) have revealed a rich set of loci and genes implicated in determining the phenotypic variation of quantitative traits. Theoretical results have also elucidated under which conditions we should expect selection response to be more or less polygenic. Is has been shown (Götsch & Bürger, 2023; Höllinger et al., 2019) that some measure of mutational target size is a crucial parameter determining the number of loci involved in adaptive architecture. Our uniquely powerful design uncovers these signatures of selection spanning an appreciable fraction of the genome. These results are consistent with the intermediate scenario from Höllinger et al. (2019), in which adaptation takes place by partial sweeps at many loci. This response is related to the strong stress caused by the high-sugar diet, as we expect several phenotypes at different levels of organization to be involved in the response to selection. While we do not have access to the phenotypes under selection, we are able to probe these putative differences by analyzing differential gene expression. Selected lines show hundreds of DE genes, and these are indeed enriched for selected SNPs, suggestive of a link between regulatory *cis*-eQTL variation and the response to selection. These widespread regulatory changes are expected to percolate through metabolic networks (Boyle et al., 2017), leading to interactions between connected genes, which potentially complicates the adaptive architecture (Forsberg et al., 2017).

We attempt to further characterize the adaptive architecture by searching for possible signatures of epistatic interactions between the SNPs under selection in the HS populations. Epistasis can be a store of additive genetic variation (Cheverud & Routman, 1995, 1996), but the effect of epistatic interaction on the long-term response to selection is thought to be modest in general (Paixão & Barton, 2016). Epistasis can become important if it is directional, i.e., if epistatic effects consistently produce positive of negative effects on the selected traits, thus enhancing or buffering additive variation (Barton, 2017; Hansen, 2013; Hansen & Wagner, 2001). Because measuring epistasis can be challenging, we have little information on the general pattern of directional epistasis (Le Rouzic, 2014), but it has been documented in some model organisms (Le Rouzic et al., 2023; Pavlicev et al., 2010). Furthermore, if several traits are selected simultaneously, the effects of epistasis on trait associations can be important in determining the response to selection (Jones et al., 2014; Melo & Marroig, 2015; Pavlicev et al., 2011). In E&R experiments, allelic redundancy has been implicated in the lack of parallelism across replicas, as segregating alleles at several loci could be combined in different ways to produce similar fitness and selective responses (Barghi et al., 2019, 2020). Under stabilizing selection, this type of interaction would necessarily lead to fitness epistasis (Bank, 2022; Höllinger et al., 2019). Alternatively, epistatic interactions can also increase parallelism, as the possible adaptive paths through genotype space are limited by the genomic background (Bank, 2022; Das et al., 2020; Kryazhimskiy et al., 2014). Given these numerous mechanisms for epistasis to contribute to long-term selection response, we searched for genomic signatures of epistatic interactions in the form of correlated allele frequencies and gametic disequilibrium across unlinked loci that where exclusive to the selected populations. We find over one thousand pairs of unlinked loci that show significantly correlated allele frequency trajectories and gametic disequilibrium exclusively in the HS populations. Using more stringent criteria, we find these signals across 11 pairwise regions, replicated in all selected populations and supported by several SNPs. While we cannot pinpoint the mechanistic source of the epistatic interaction we observe, we can detect them with high confidence, indicating that there is a potential role for epistasis in the adaptive architecture for high-sugar stress that should be further explored.

## Methods

### Mapping Population

To allow the detection of allelic effects that would be hidden in natural populations due to low frequency, we created a synthetic outbred mapping population. To create this population, we selected 16 inbred lines from the Netherlands population (NEX) from the GLOBAL DIVERSITY LINES (Grenier et al., 2015). The lines were selected based upon their low frequency of inversions to reduce the suppression of recombination associated with inversions (Barghi & Schlötterer, 2019). To establish the population from these lines, we performed a round-robin cross on the initial lines (1 × 2, 2 × 3, …, 16 × 1) and subsequently performed a round-robin cross on the F1s to ensure parental representation and that no chromosome was lost. The resulting F2 individuals were placed in a x b x c cm cages and allowed to recombine freely for more than 50 generations. This design increases the allele frequency of rare variants by replicating and randomizing throughout the population (fig. 1 C).

### Selection Regime

We performed a laboratory natural selection experiment (Fuller et al., 2005) on high-sugar diets without selecting for any phenotype. High-sugar diets are known to have high fitness costs (Musselman et al., 2011; Na et al., 2013; Pallares, Lea, et al., 2020) and by allowing our populations to directly evolve under this physiological stress, we explored the adaptation to this deleterious effect. To do this, we subdivided our mapping population into 6 replicate populations, 3 of which were placed on a standard medium and 3 of which were placed on highsugar medium. The standard medium consists of 8% glucose, 8% yeast, 1.2% agar, 0.04% phosphoric acid, and 0.4% propionic acid. High-sugar medium follows the same recipe as standard medium with the addition of 12% glucose resulting in a total of 20% glucose. Each population was placed in a population cage (BugDorm #4F3030) and maintained at ∼5000 individuals for ∼120 generations. Each generation was seeded from an egg lay, on fresh bottles of the respective diet, at 5-6 days post-eclosion. After pupation but before eclosion, bottles were cleared of adults, moved to new cages and opened. Following each egg lay, individuals were collected and stored at -80 C for subsequent sequencing.

### Factorial egg lay after selection

To assess if selected populations had adapted to the stressful high-sugar environment, we performed a factorial egg lay experiment, measuring the fecundity of both CONTROL and HS populations in the control and highsugar diet. We started by controlling/standardizing population density for the parental flies. To that end, adult flies from each selection regime (control and HS) laid eggs on apple juice-agar plates for 3 hours. These plates were then incubated at 24 degree C, 70% relative humidity, and 12:12 Light/Dark cycle for 24hrs to obtain 1st instar larvae. We then placed 50 larvae (low density) in each vials with 8-10ml of the appropriate food type (control or high sugar), the emerging flies were used for the fertility measurement. At the time of the assay, all flies were between 4-6 days old. Flies from each treatment were allowed to lay eggs on a molasses-agar medium with a small amount of yeast paste. These plates were switched every 2 hrs, 3 times on one day and the same process was repeated the following day. Eggs on each of the plates were counted to provide a fecundity estimate. Both populations show higher fitness in the diet to which they had adapted (fig. 1 D).

### Library preparation and sequencing

Flies from generation 1, 11, 25, and 100 were selected from each population for sequencing and distributed into 96 well plates. One 2.8 mm stainless steel grinding bead (OPS diagnostics, #089-5000-11) and 100 µl of lysis buffer were added to each well. Flies were homogenized for 10 minutes at maximum speed in a Talboys High Throughput Homogenizer (#930145). The resulting lysate was moved to a new 96-well plate for DNA extraction, using a Multi-Well Plate Vacuum Manifold (Pall Life Sciences #5017) and Acroprep advance 1 ml DNA binding plates (Pall Life Sciences #8132).

Library prep was performed using a liquid handling robot (CyBio® FeliX, Analitik Jena) to ease the processing of many samples and reduce variability from manual handling of samples. The protocol broadly followed the strategy described in Picelli et al (Picelli et al., 2014). Specifically, we added 10 *µ*l (100 *µ*M) of forward oligo adapter A and 10 *µ*l (100 *µ*M) of reverse oligo adapter (Tn5MERev) to 80 *µ*l of reassociation buffer (10 mM Tris pH 8.0, 50 mM NaCl, 1 mM EDTA). Following this, we annealed in a thermocycler with the following program: 95°C for 10 minutes, 90°C for 1 minute, reduce the temperature by 1°C per cycle for 60 cycles, and then hold at 4°C. The process was repeated for oligo adapter B. To load the adapters onto Tn5, we mixed 5 *µ*l of Tn5, 9 *µ*l of pre-annealed adapter A, and 9 *µ*l of pre-annealed adapter B then incubated this mixture in a thermocycler at 37°C for 30 minutes. The resulting pre-charged Tn5 was the diluted with a 1:1 solution of reassociation buffer and glycerol to 1:1 reassociation buffer:glycerol to pre-charged Tn5.

### Mapping of reads, SNP calling, and estimation of allele frequencies

Following sequencing, we mapped reads to the *Drosophila melanogaster* reference genome (v6.14) using BWA (3) (Li & Durbin, 2009), retained only uniquely mapped reads, and removed PCR generated duplicates using Picard (“Picard Toolkit,” 2019). SNPs were called jointly in batch 1 (generations 1,11,25; 1728 samples) and batch 2 (generation 100; 1116 samples) using the haplotype-based variant detector Freebayes (Garrison & Marth, 2012), ignoring indels and multi-allelic SNPs. Any SNPs with a quality score less than 30, or with a coverage smaller than 28x in any population in batch 1, or smaller than 80x in any population in batch 2, were excluded. We also excluded SNPs with coverage above the genome wide baseline of 167x in any population in batch 1, or above 333x in any population in batch 2, since such highly covered SNPs might be indicative of collapsed repeats. After these filtering steps, we retained 1,741,428 SNPs on the major chromosomes (2L, 2R, 3L, 3R and X) for subsequent analyzes.

Allele frequencies inferred from pooled sequencing can be biased if the coverage per individual in the pool is uneven (Schlötterer et al., 2014). Our individually barcoded DNA-libraries allowed us to identify from which individual any given read originates, thereby avoiding this problem. The variance in per sample read depth was substantial, suggesting that allele frequency estimated from Pool-seq, agnostic to read origin, might be error prone. We corrected for uneven coverage when estimating allele frequencies using the formula: 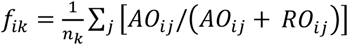, where *f*_*ik*_ is the estimated allele frequency of SNP *i* in pool *k, n* is the number of individuals in the pool, and *AO*_*ij*_ and *RO*_*ij*_ are the number of observations of the alternative and reference alleles respectively at SNP *i* in individual *j*. The sum is taken over all individuals in pool *k*, where *k* is one of the 24 [4 (timepoints) x 3 (replicate populations) x 2 (treatments)] population samples. This strategy corrects for unequal coverage and should be refered over naive Pool-seq estimates. We thus obtained estimates of allele frequency per SNP, in each of the 24 population samples.

We minimized variability in sequence coverage introduced by manual handling of samples by preparing libraries with a CyBio® FeliX liquid handling robot. We pooled libraries according to DNA concentration, and re-pooled the DNA libraries after preliminary sequencing on the Illumina Miseq platform to normalize coverage.

### Inference of patterns of polygenic adaptation using PCA

To explore the main drivers of genetic changes during the course of the experiment, we performed a principal component analysis of the genome wide allele frequencies. For this analysis, we first assembled a sample-by-SNP matrix P, containing the genome-wide allele frequencies in all 24 population samples (4 time points x 2 treatments x 3 replicate populations). We then computed the principal components of this matrix, allowing us to identify the main drivers of allele frequency changes in an unsupervised fashion.

### Inference of individual loci selection signatures

To detect signals of positive selection, we fitted the following logistic regression model:

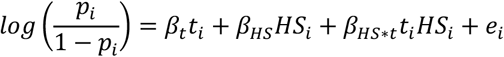

where *p*_*i*_ denotes the allele frequency at a given locus in a given population and time point; *t*_*i*_ is a numerical variable corresponding to the 4 sampled time-points (generations 1, 11, 25, and 100 are numerically coded as }1, 2, 3, 4}, and so the corresponding coefficient (*β*_*t*_) measures the average allele frequency change across all time-points); *HS*_*i*_ is an indicator variable corresponding to the two treatments *HS* = 1 in high sugar (HS) and *HS* = 0 in control (CONTROL)); *e*_*i*_ is an error term. The β parameters are the corresponding regression coefficients. This allowed us to model the allele frequency for every locus across the entire selection experiment in one joint statistical framework. We focus primarily on the interaction effect β_HS*t_, which quantifies the degree to which the allele frequency trajectory in the control regime differs from the one in the high-sugar regime.

After fitting this model for all SNPs, we obtained estimates of the effect of time separately for the control and highsugar selection regimes. This was done using the emtrends function in the R package *emmeans* (Lenth, 2022). In order to exclude selection signatures that did not correspond to high-sugar adaptation, we disregarded SNPs where the effect of time in the high-sugar selection regime showed a p-value above 10^−4.^

### Individual level genotypes

To obtain individual-level genotypes rather than allele frequencies from our low coverage data at generation 100, we first filtered our SNP data more stringently. Having already applied the filter described above, we retained SNPs with called genotypes in more than 90% of the individuals, each genotype being called with a minimum depth of 3. We also excluded individuals with more than 50% missing genotypes. This filtering was done separately for each chromosome, giving a set of 51k SNPs called in 412 individuals in the control populations, and 52k SNPs called in 439 individuals in the high-sugar populations. These genotypes where used to estimate linkage disequilibrium (fig. 1 B) and to search for signatures of selective sweeps and gametic disequilibrium.

### Detecting selective sweeps

To detect signatures of selective sweeps, we first used the software shapeit (Delaneau et al., 2008) to phase the individual genotypes into haplotypes. The estimated haplotypes were then used to calculate the integrated Haplotype Score (iHS) (Voight et al., 2006). Briefly, iHS measures the length of haplotype homozygosity around a given allele, compared to its alternative allele. A recent selective sweep is expected to leave a genomic footprint of extended homozygosity around the selected allele, whereas selection on standing genetic variation and/or polygenic selection might not leave such a footprint (Lynch & Walsh, 1998). iHS was calculated using the Rpackage rehh (Gautier & Vitalis, 2012), and scores were standardized per allele frequency bin as described in Voight et al. (2006). We calculated iHS at generation 100 in the high-sugar selected populations, using the 3 replicate populations. To compare selection signatures inferred from our regression model to selective sweeps inferred by iHS, we contrasted the regression p-values to iHS on a SNP-by-SNP basis (sup fig. 3).

### Transcriptional changes associated with adaptation to high-sugar diet

#### Experimental design

To identify transcriptional changes associated with genetic adaptation to high sugar, we performed an experiment that allowed us to robustly differentiate gene expression differences due to the adaptation regime from the plastic response due to short-term changes in dietary condition. For this, we used a full reciprocal design where flies from each replicate cage from generation 170 were allowed to lay eggs in either the dietary condition they evolved in (i.e., HS evolved flies on high-sugar food, CONTROL evolved flies on control food), or in the alternative diet (i.e., HS evolved flies on control diet, CONTROL evolved flies on high-sugar diet). Female flies were collected 7-11 days after eclosion, and head and body were separated and plated each in two 96-well plates with each plate containing samples for only one tissue and all four experimental combinations. Plates were stored at -80°C until further processing.

#### RNA extraction and sequencing

Plates containing heads and bodies were processed in the same way: Sample homogenization was done as described above for DNA samples, and mRNA extraction as described in Suppl. File 2 of Pallares, Picard, et al. (2020) using Dynabeads™ mRNA DIRECT™ Purification kit (ThermoFisher), and a final elution of 10 *µ*l and 30 *µ*l Tris-HCl for heads and body, respectively. 3’-enriched RNAseq libraries were prepared following the TM3’seq pipeline (Pallares, Picard, et al., 2020). In brief, 10 *µ*l of input mRNA was used in the first strand cDNA synthesis reaction which was primed with Tn5Me-B-30T oligo that binds to the polyA tail of mRNA molecules resulting in 3’ enriched libraries. cDNA was amplified in three rounds of PCR and tagmented using homemade Tn5 transposase. 12 PCR cycles were used for final library amplification using Illumina’s i5 and i7 primers. The step by stepTM3’seq protocol can be found in Suppl. File 1 of Pallares, Picard, et al. (2020). All libraries within a plate were pooled using 5 *µ*l or 2 *µ*l per head and body library, respectively, and cleaned and size-selected using the double-sided Agencourt AMPure XP bead (Beckman Coulter) cleanup approach described for DNA-seq libraries. The resulting four plate-level libraries were pooled in equal proportions and sequenced on the Illumina NovaSeq S2 platform at the Genomics Core Facility of the Lewis-Sigler Institute for Integrative Genomics at Princeton University. RNA extraction, cDNA synthesis, and library preparation were done in the CyBio® FeliX liquid handling robot.

#### Processing of RNAseq data

Raw RNA-seq reads were trimmed to remove low quality bases, adapter sequences, and to exclude post-trimmed reads shorter than 20 nt using Trimmomatic 0.32 (Bolger et al., 2014) and the following parameters: SE ILLUMINACLIP:1:30:7 LEADING:3 TRAILING:3 SLIDINGWINDOW:4:15 MINLEN:20. The trimmed reads were mapped to the *Drosophila melanogaster* genome r6.14 using STAR (Dobin et al., 2013), and uniquely mapped reads were assigned to genes using feautureCounts from the package Subread (Liao et al., 2013) and the following parameters: -t exon –g gene_id]. Samples with fewer than 500k or more than 20M gene counts, and genes with mean CPM < 1 were removed. After this filtering, the final dataset used in further analysis consisted of 161 head samples with a median of 3.45M gene counts covering 8460 genes, and 171 body samples with a median of 2.3M gene counts covering 8360 genes.

#### Differential expression analysis

To identify the transcriptional differences due to adaptation to high sugar, we performed a differential expression analysis between flies evolved in HS diet and flies evolved in CONTROL diet while accounting for the dietary condition the flies were exposed to for one generation. For each tissue separately, we used a Wald test in DESeq2 (Love et al., 2014) and the following design: Expression ∼ Plate + Diet + Genotype, where Plate indicates the 96-well plate in which samples were processed from sample collection through library preparation; Diet represents the dietary condition the flies were exposed to for one generation (HS or CONTROL); Genotype represents the diet flies evolved in (HS or CONTROL). Sample size for each of the four groups in body and head, respectively: *n*(genotype HS, diet HS) = 41, 38; n(genotype HS, diet CONTROL) = 46, 42; *n*(genotype CONTROL, diet CONTROL) = 45, 41; *n*(genotype CONTROL, diet HS) = 39, 40. p-values were estimated for the null hypothesis lfcThreshold = 0 and alpha = 0.05, and adjusted using Benjamini & Hochberg FDR method.

#### Differentially expressed genes and selection

Having identified genes that were differentially expressed between flies adapted to the respective selection regimes, we went on to look for signals of selection nearby along the chromosome to these genes. Considering an interval ± 5 kb around each gene, we looked for SNPs showing a significant time-by-regime effect in the regression analysis and overlapped with the DE genes. To this end we used the R-package GenomicRanges, and the fraction of overlapping selected SNPs was calculated at p-value thresholds }10^−5^, 10^−10^, 10^−15^, 10^−20^, 10^−25^, 10^−30^} for the time-by-regime effect (fig. 5 B, red lines and points). Because of the large number of DE genes and selected SNPs, we expect some amount of overlap between the two by chance. To quantify this expected chance overlap, we performed a permutation test. For each permutation, the same number of SNPs as we observed to be significant at that significance threshold was picked at random from the full set of 1.76M SNPs. We then calculated the fraction of these random SNPs that overlapped with the DE genes. Performing 1000 permutations at each p-value threshold gave us empirical null distributions for the overlap (fig. 5 B, boxplots).

### Epistatic selection signatures

#### Wright-Fisher model with selection for epistatic QTLs

We used an individual based Wright-Fisher model to investigate the effect of epistatic interactions in the interchromosomal LD in our selection experiment. The simulation was based on the code from Lou et al. (2020), using the SLiM modeling framework. We start by creating a neutral burn-in population with 5000 individuals, two equal chromosomes with 300 k sites, a base mutation rate of 1.5×10^−9^, and a between-site recombination rate of 10^−8^. This burn-in population is allowed to evolve under a Wright-Fisher neutral model for 50k generations. With these parameters we expect about 0.5 recombinations per generation, and after 50k generations we have about 3k segregating neutral SNPs in mutation-drift equilibrium with a minor allele frequency above 5%. The burn-in process was repeated for each simulation replicate, so each replicate simulation started with a different initial population. For each initial population, we also did experimental replicates, which started from the same starting population.

Using these starting populations, we sampled 1000 of the segregating mutations to be QTLs contributing to the phenotypic effect of a polygenic trait, with all QTLs having the same phenotypic effect and 2 alleles. We then considered two scenarios: (1) an additive scenario, where the value of the trait is only given by these additive QTLs, and (2) an epistatic scenario, where, in addition to the additive effects, we sampled 200 pairs of QTLs (one member of the pair in each chromosome) to have an additive-by-additive epistatic effect. In this epistatic scenario, the value of the trait depended on these epistatic interactions. Both scenarios proceed with truncation selection on the polygenic trait for 100 generations, with the 500 individuals with the smallest trait value being removed before reproduction in each generation. This strength of selection was chosen so that at the end of the simulation we had only a few fixations (around 10), mimicking the observation in our fly selection experiment.

During the selection phase of the simulation we sampled allele frequencies at regular intervals (every 10 generations) and used these to calculate the correlation between allele frequencies at the QTL pairs. At generation 100, we also measured the gametic disequilibrium between the same pairs of QTLs in both simulations, with the only difference being the presence of the epistatic interaction in one of the scenarios. We compared the mean gametic disequilibrium between these QTL pairs to a distribution of the mean gametic disequilibrium between random pairs of SNPs across chromosomes. To create this distribution, we sampled 200 SNPs in each chromosome and calculated the gametic disequilibrium between the pairs, this process was repeated 10k times. We created a separate distribution for each scenario.

#### Identifying well-supported epistatic SNP pairs

To detect potential instances of epistatic selection, we looked for two types of signals: 1) correlations between the allele frequencies at different loci; 2) gametic disequilibrium after 100 generations of selection. The former could be due to epistasis or due to similar but independent selection coefficients at the respective loci, while the latter is only expected under epistasis. The correlations were estimated by the Pearson correlation coefficient, using allele frequencies in all generations in the high-sugar populations (*n*= 12 per SNP pair). The gametic disequilibrium was quantified separately in highsugar and control populations, using the individual genotypes at generation 100 described above. For each SNP pair, we tested the deviation from independent segregation using a chi square test (*n*∼ 420 per SNP pair). Having identified candidate pairs where the two SNPs displayed gametic disequilibrium and were located on different chromosomes, we attempted to find well supported signals by clustering physically close SNPs with a similar signal of gametic disequilibrium. To do this, we applied a clustering procedure akin to the LD clumping algorithm implemented in PLINK (Purcell et al., 2007). The algorithm works as follows:

1. Starting with the SNP pair with the smallest p-value for gametic disequilibrium, assign to the same cluster all other SNP pairs with one SNP on chromosome 2 that are within 250 kb, in linkage disequilibrium, and have not already been clustered. Repeat until there are no more SNP-pairs to assign to clusters. Thus, each cluster contains SNP pairs sharing proximal and linked SNPs on chromosome 2.
2. For each cluster identified in (i), perform a second round of clustering for the SNP pairs within that cluster. This is done by assigning to the same cluster all SNP-pairs with one SNP on chromosome 3 that are within 250 kb, in linkage disequilibrium, and have not already been clustered.

We thus obtain hierarchical clusters, where each “chromosome 2 end” cluster contains one or several “chromosome 3 end” clusters. The algorithm is greedy, so each SNP pair will only end up in one cluster, if at all. Finally, we keep the clusters with at least 3 linked SNPs at “each end”. We thus identify epistatic selection signatures supported by multiple SNP pairs, with multiple linked SNPs at each end of the putative interaction. Through the nature of our clustering procedure, each such signature will involve one locus on chromosome 2, and one or several loci on chromosome 3 (fig. 6 B).

## Code and data availability

Code to reproduce figures in this paper is available at github.com/ayroles-lab/highsugar-selection-code.

Corresponding data is available at **Princeton DataSpace link**.

## Supporting information and figures

Supporting information can be found at github.com/diogro/HighSugarSelection. response. *EMBO J*., *21*(22), 6162–6173. https://doi.org/10.1093/emboj/cdf600

**Supplementary Figure 1:**
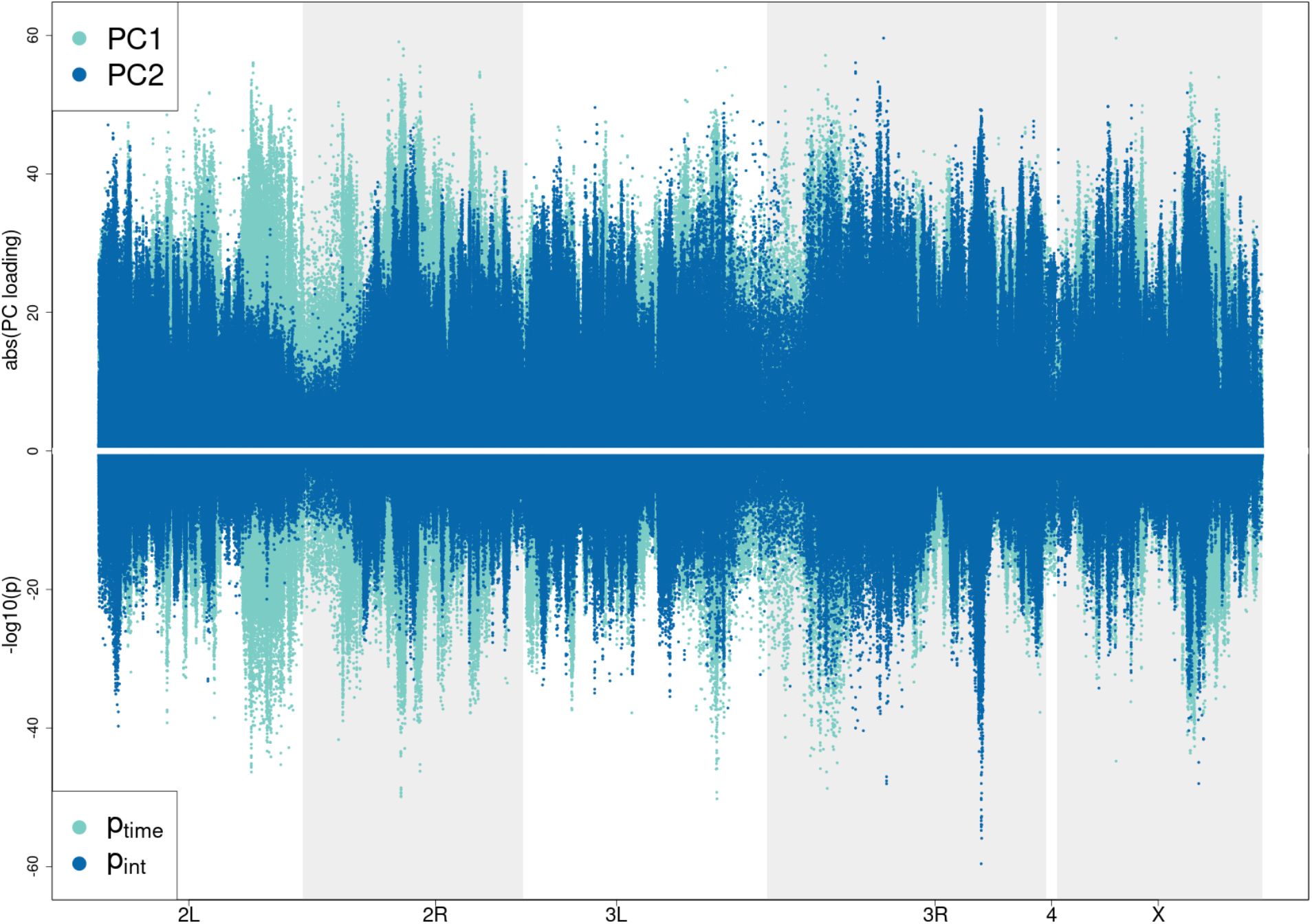
Comparison of PC scores and p-values from regression analysis. **PC1** scores are similar to p-values from SNPs that show a shared selection signal across treatment and control, and PC2 scores are similar to p-values from SNPs that only show selection signal in HS. These similarities corroborate the interpretation that PC1 is related to the SNPs involved in the shared selection captured by the time component of the regression, and that PC2 is related to the SNPs involved in the response to high-sugar selection, which is captures by the time-by-selection regime interaction coefficient in the regression analysis.

**Supplementary Figure 2:**
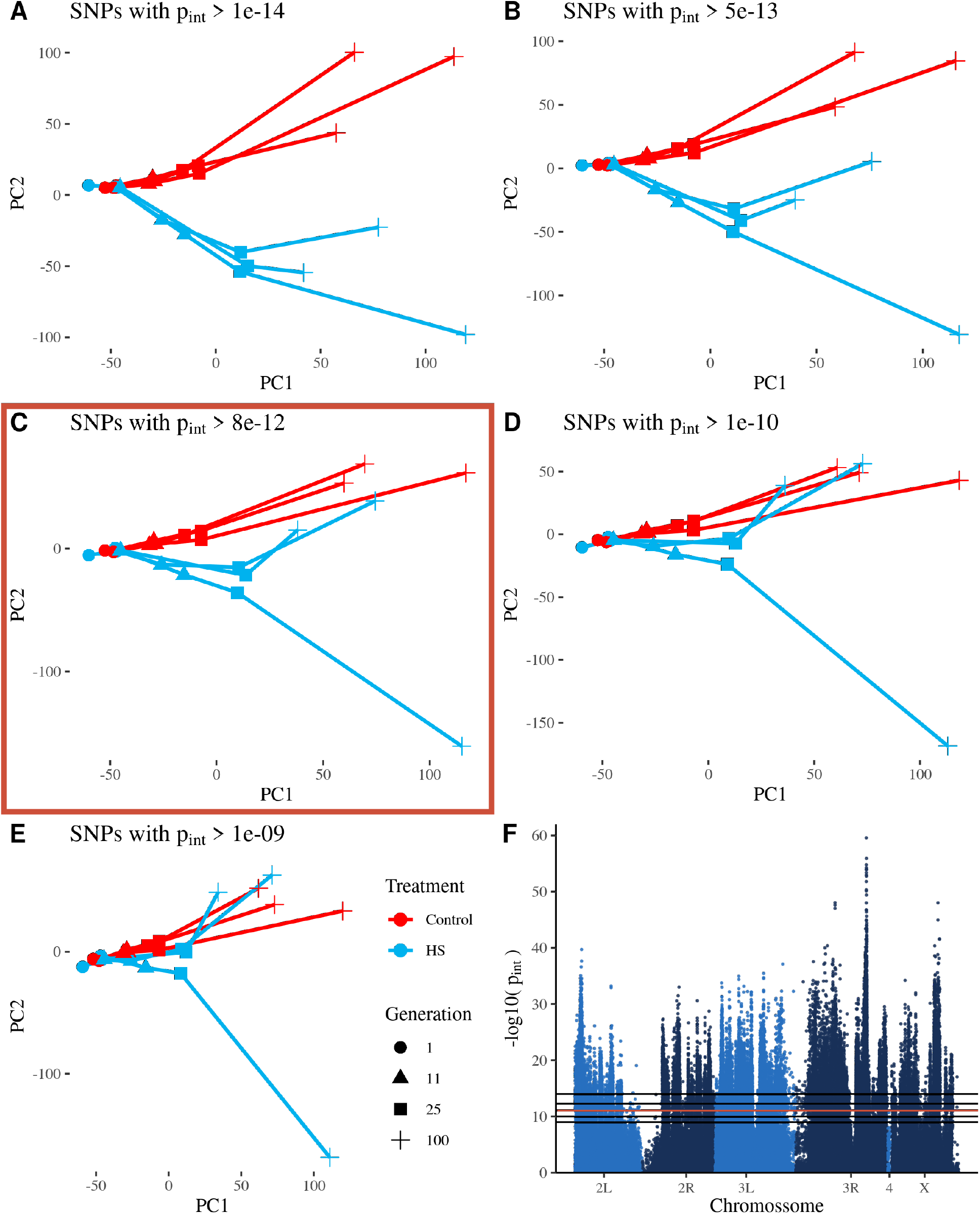
Across population allele frequency PCA (as in fig. 2) after exclusion of significant SNPs. As the p-value significance threshold for the high-sugar selection interaction term is increased and more putatively selected SNPs are removed, PC2 gradually explains less and less of the divergence between HS and CONTROL populations. The Manhattan plot in the final panel shows the -log10 p-values for the interaction term in the regression analysis and the horizontal lines show the various thresholds used in the PCA panels. For each PCA panel, SNPs above the threshold are removed before calculating the PCA. The threshold we chose based on the differentiation between CONTROL and HS populations in the PCA plot is marked by the red line in the Manhattan plot and the red box around the corresponding PCA plot.

**Supplementary Figure 3:**
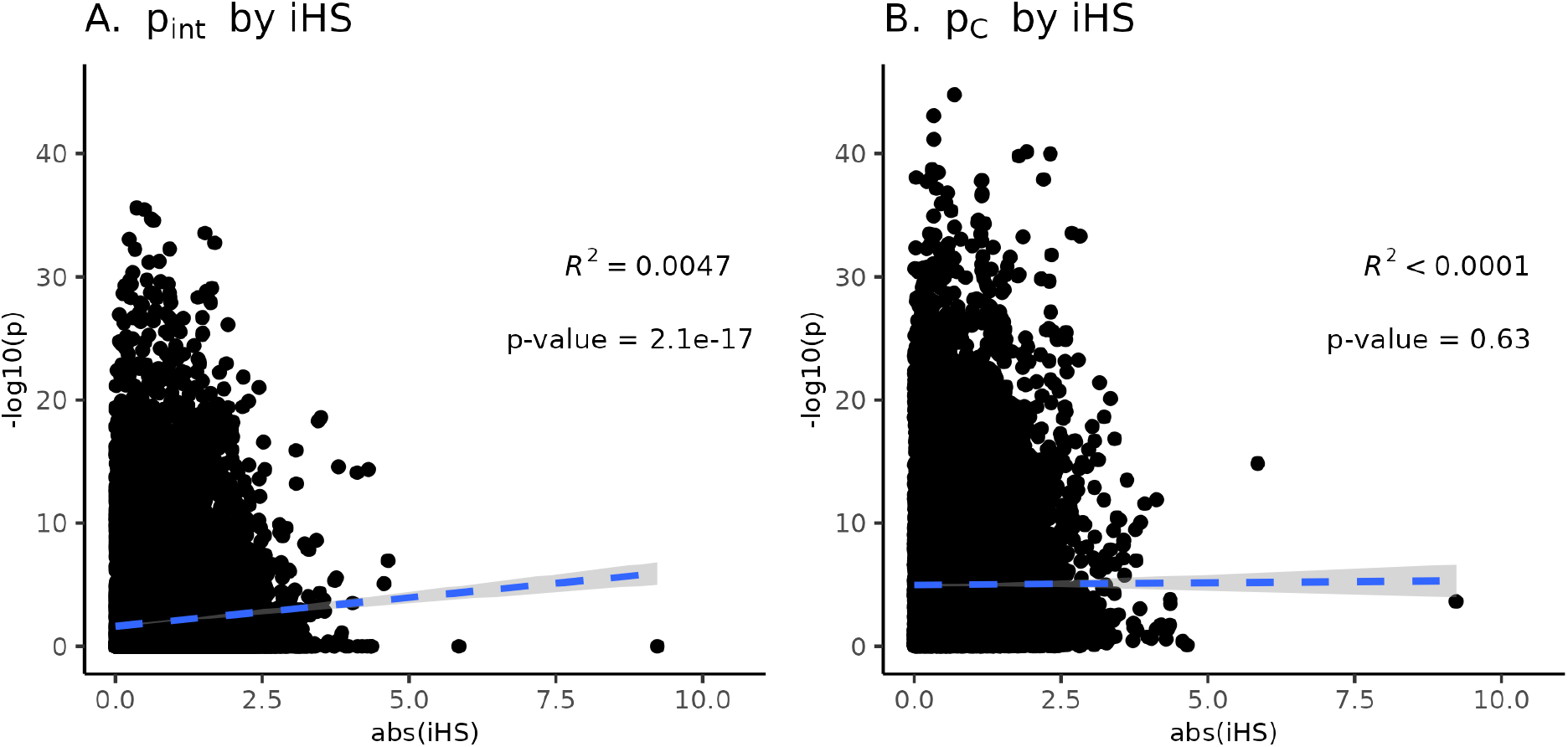
A comparison of the results from the time-series regression analysis and the iHS analysis. We compare the p-values from the regression analysis to the absolute iHS values calculated on the HS populations. (A) Negative log10 transformed p-values corresponding to the time-by-selection regime interaction coefficient in the regression analysis of allele frequency over time (y-axis) plotted versus absolute iHS scores (x-axis). (B) Negative log10 transformed p-values corresponding to the CONTROL population contrast in the regression analysis of allele frequency over time (y-axis) plotted versus absolute iHS scores calculated on the HS populations (x-axis). Around 15k SNPs included in the figure, with each point representing one SNP. Only the interaction term p-value are associated with iHS values, suggesting both capture a signal related to high-sugar selection. In contrast, signals associated with changes in the CONTROL population are not associated with the iHS signal in the HS populations.

